# Decoding the chromatin proteome of a single genomic locus by DNA sequencing

**DOI:** 10.1101/229955

**Authors:** Tessy Korthout, Deepani W. Poramba-Liyanage, Kitty F. Verzijlbergen, Tibor van Welsem, Fred van Leeuwen

## Abstract

Transcription, replication and repair involve interactions of specific genomic loci with many different proteins. How these interactions are orchestrated at any given location and under changing cellular conditions is largely unknown because systematically measuring protein-DNA interactions at a specific locus in the genome is challenging. To address this problem, we developed Epi-Decoder, a Tag-ChIP-Barcode-Seq technology in budding yeast to identify and quantify in an unbiased and systematic manner the proteome of an individual genomic locus. Epi-Decoder is orthogonal to proteomics approaches because it does not rely on mass spectrometry but instead takes advantage of DNA sequencing. Analysis of the proteome of a transcribed locus proximal to an origin of replication revealed more than 400 proteins. Moreover, replication stress induced changes in local chromatin-proteome composition prior to local origin firing, affecting replication proteins as well as transcription proteins. Epi-Decoder will enable the delineation of complex and dynamic protein-DNA interactions across many regions of the genome.

## Introduction

The chromatin at any given location in the genome is a dynamic entity that likely involves the interaction of many different proteins and non-protein factors. The essential and complex processes of transcription, replication and repair require major chromatin rearrangements to access and use the genome^1-2^. Furthermore, the genome's chromatin is under the influence of signals that can relay information of cellular events or states to the genome and vice versa^3-4^. Fully understanding the chromatin-regulatory mechanisms in the cell will require comprehensive knowledge of the full set of proteins that bind at individual genomic loci. Although many chromatin factors have already been identified by genetics and protein-protein interaction studies, direct, unbiased and comprehensive analyses of chromatin interactions at specific genomic loci has remained a major challenge^5-6^. A commonly considered strategy towards solving this problem is introducing an affinity handle at a locus of interest, purifying the locus (capture) and analyzing the co-purified proteins by mass spectrometry (MS)^7-8^. However, a major challenge with capture-MS is the need for very high levels of enrichment of the locus of interest versus the rest of the large genome while at the same time obtaining sufficient amounts of material for comprehensive and quantitative MS analysis^5-6^. For example, in a model organism with a small genome such as yeast, purification of a 1 kb locus with its associated proteins from an entire genome of ~30 Mb requires a 30,000-fold purification.

Here, we present an independent approach, Epi-Decoder, with which the interactome of a single-copy locus is determined by DNA sequencing instead of mass spectrometry. In this approach developed in yeast, a library of clones is generated in which each clone harbors at least one DNA barcode at a fixed locus and one protein tagged at its endogenous locus with a common epitope tag. Following chromatin immunoprecipitation (ChIP) for the common tag on a pool of cells, the barcodes of the co-immunoprecipitated DNA are counted by high-throughput sequencing and the amount of barcode recovered serves as a read-out for the amount of the corresponding specific protein binding at the barcoded locus.

Epi-Decoder of a single transcribed locus in yeast identified more than 400 chromatin-interacting proteins, enabled the identification of differences in protein binding between the 5' and 3' end of a gene, and demonstrated a chromatin rewiring in response to physiological changes. Thus, Epi-Decoder provides an efficient method to provide a comprehensive map of the dynamic proteome of a single-copy locus. Moreover, it is an orthogonal approach to capture-MS because quantitative and qualitative information of protein binding does not involve mass spectrometry but is obtained by DNA sequencing. Epi-Decoder should be widely applicable to many genomic loci and will greatly facilitate the delineation of complex and dynamic protein-DNA interactions of the genome.

## Results

### Epi-Decoder: Measuring protein binding by DNA barcode counting

To develop a strategy for systematic and comprehensive decoding of the interactome of a single genomic locus, we asked whether DNA barcode technology can be harnessed to solve this challenging proteomics problem. We and others previously showed that short DNA barcodes (< 20 bp) integrated in the genome and embedded in chromatin can serve as molecular identifiers of the chromatin state they are in^9-11^. Building on that notion, we used Synthetic Genetic Array (SGA) technologies^12-13^ to create arrayed yeast libraries that allow for high-throughput and direct assessment of chromatin states of many cell clones in parallel. In these libraries, each clone contains a pair of known unique barcodes flanking a constitutively transcribed kanR marker gene under control of the Ashbya gossypii *TEF1* promoter and terminator at the *HO* locus^14^ as well as a known Tandem Affinity Purification (TAP)-tagged protein expressed from its endogenous locus^15^ (Fig. 1a). The Epi-Decoder libraries used here cover ~4000 TAP-tagged proteins (Supplementary Table 1) of the ~5600 proteins encoded in the yeast genome^16^. Following pooling of the barcoded TAP-tag strains, cells are incubated with formaldehyde to crosslink proteins to DNA and subjected to ChIP (Fig. 1b). From the co-immunoprecipitated DNA and the input samples, the barcoded regions are amplified (Supplementary Fig. 1), and the barcodes identified and quantified by parallel sequencing (Fig. 1b). In this set-up, which we call Epi-Decoder, the abundance of a barcode (ChIP/input) reports on the cross-linking of the corresponding TAP-tagged protein to its own barcoded region for every TAP-tagged clone in the pool (Fig. 1c). The reporter gene at the *HO* locus contains two barcodes that can be analyzed in parallel but have different chromatin contexts. The upstream barcode (BC_UP) is located in the promoter region whereas the downstream barcode (BC_DN) lies in the terminator region and in close proximity to an origin of replication (Fig. 1c). Thus, Epi-Decoder results in an inferred binding score for the vast majority of the proteins present in the yeast genome at two distinct genomic loci.

**Figure 1.**
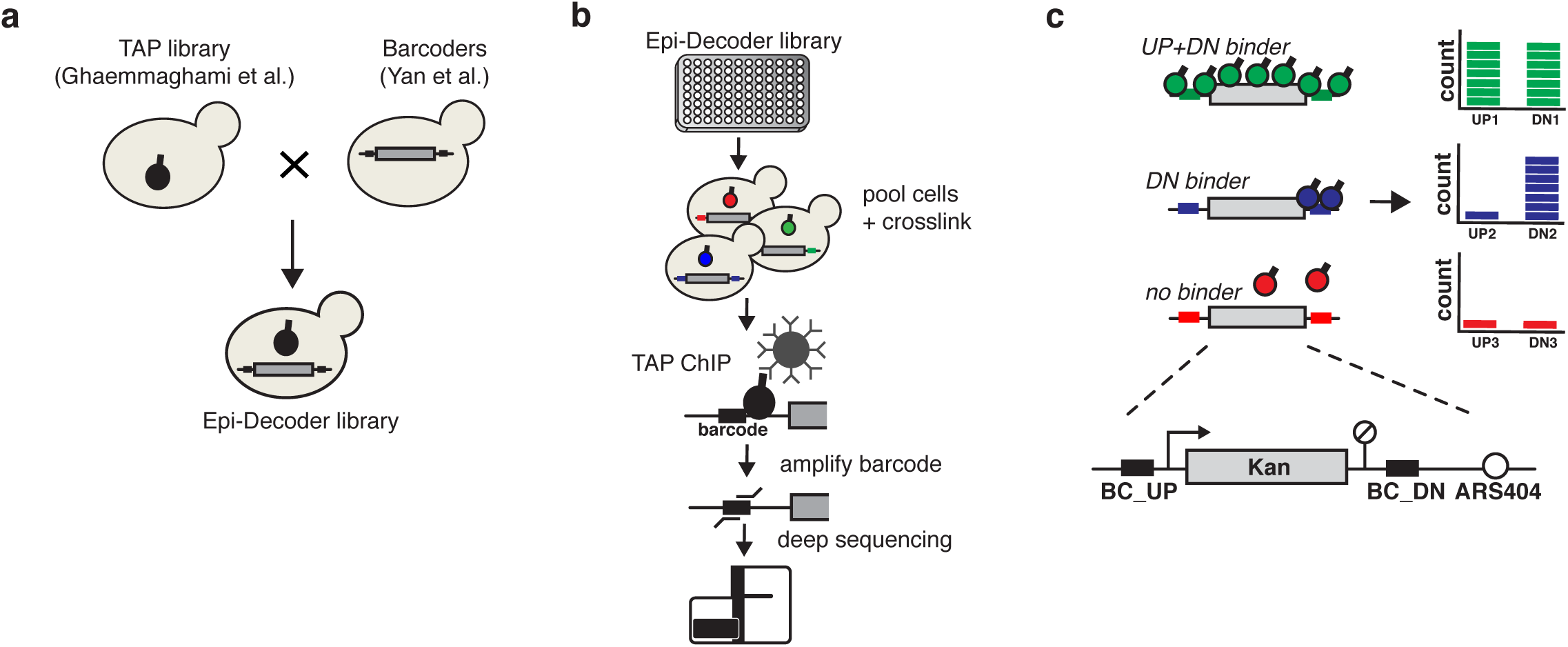
Outline of Epi-Decoder. (A) Construction the Epi-Decoder library: a library of strains encoding TAP-tagged proteins was crossed to a Barcoder library by using standard SGA procedures. (B) Experimental set-up of Epi-Decoder: colonies of an arrayed library are pooled in a flask, expanded, subjected to cross-linking, and used for a TAP ChIP. The co-immunoprecipitated barcodes are amplified and counted by massive parallel sequencing. Barcodes of input DNA are counted to control for the presence of each barcode in the pool. Indexed input and ChIP samples of multiple pools can be combined into one master pool before sequencing. (C) Protein binding is measured by barcode counting. The Barcoder locus consists of a KanMX resistance marker transcribed by the *TEF1* promoter flanked by two unique barcodes and replacing the HO gene. The BC_UP is promoter-proximal, the BC_DN terminator-proximal and next to an origin of replication (ARS404).

### Epi-Decoder identifies many chromatin-associated factors

For BC_UP and BC_DN we identified the immunoprecipitated barcodes with significantly different counts compared to background (red dots in Fig. 2a), based on six biological replicates (Supplementary Fig. 2a). The vast majority of significantly different barcodes showed a positive binding score (Fig. 2a, red dots, IP/Input > 0). Here we refer to the factors associated with those barcodes as binders. Together, we identified 469 binders of which 18 were specific for BC_UP and 273 for BC_DN (Fig. 2b, Supplementary Table 1). Significantly depleted factors (Fig. 2a, red dots IP/input<0) were not expected because proteins cannot bind less to their barcodes than negative controls such as non-expressed or non-tagged proteins. The low number of significantly depleted factors reflect false discoveries, in agreement with our stringent cut-off (FDR<0.01). Among the binders, the four canonical histone proteins (represented by seven histone genes in the library) were among the most-enriched proteins at both barcodes. Furthermore, 267 out of the 469 binders had GO terms related to DNA binding (Fig. 2c, Supplementary Fig. 2b), confirming that Epi-Decoder provides an effective approach for identifying chromatin-interacting factors. In addition to known DNA-interacting proteins, we found a substantial number of factors involved in RNA processing and cellular metabolism. RNA processing events such as capping, splicing, cleavage, and polyadenylation are all processes known to occur in close conjunction with RNA transcription and hence in proximity to DNA^17^, providing an explanation for the recovery of barcodes associated with RNA processing proteins. The presence of factors involved in cellular metabolism cannot be explained by general associations with transcription, although recent studies have suggested various roles of metabolic enzymes in the nucleus^4,18-19^ (and see discussion). Given the fact that some of these factors are known to be highly expressed, we investigated the possibility that their chromatin interactions were determined by protein abundance. Overall, chromatin binders were generally more highly expressed than non-binders (Fig. 2d). However, high protein expression level alone was not sufficient for binding: many chromatin binders were not highly abundant and many highly abundant proteins were non-binders. For example, ribosomal proteins are highly abundant but most of them were not detected as chromatin interactors even though ribosomal proteins traffic through the nucleus to form ribonucleoprotein complexes for ribosome assembly. Furthermore, for a selected panel of factors reflecting different classes of binders we could quantitatively confirm the positive and negative Epi-Decoder results by ChIP-qPCR analysis of individual clones (Fig. 2e). Together, this suggests that the barcode counts obtained by Epi-Decoder accurately and quantitatively report on the efficiency of protein crosslinking at the barcoded loci and captures histones, other core chromatin proteins as well as additional factors. We note that in the Epi-Decoder set-up, proximity to DNA, genomic distance from the barcode, protein abundance, and cell-to-cell variation are among the factors that can influence the measured quantitative binding scores.

**Figure 2.**
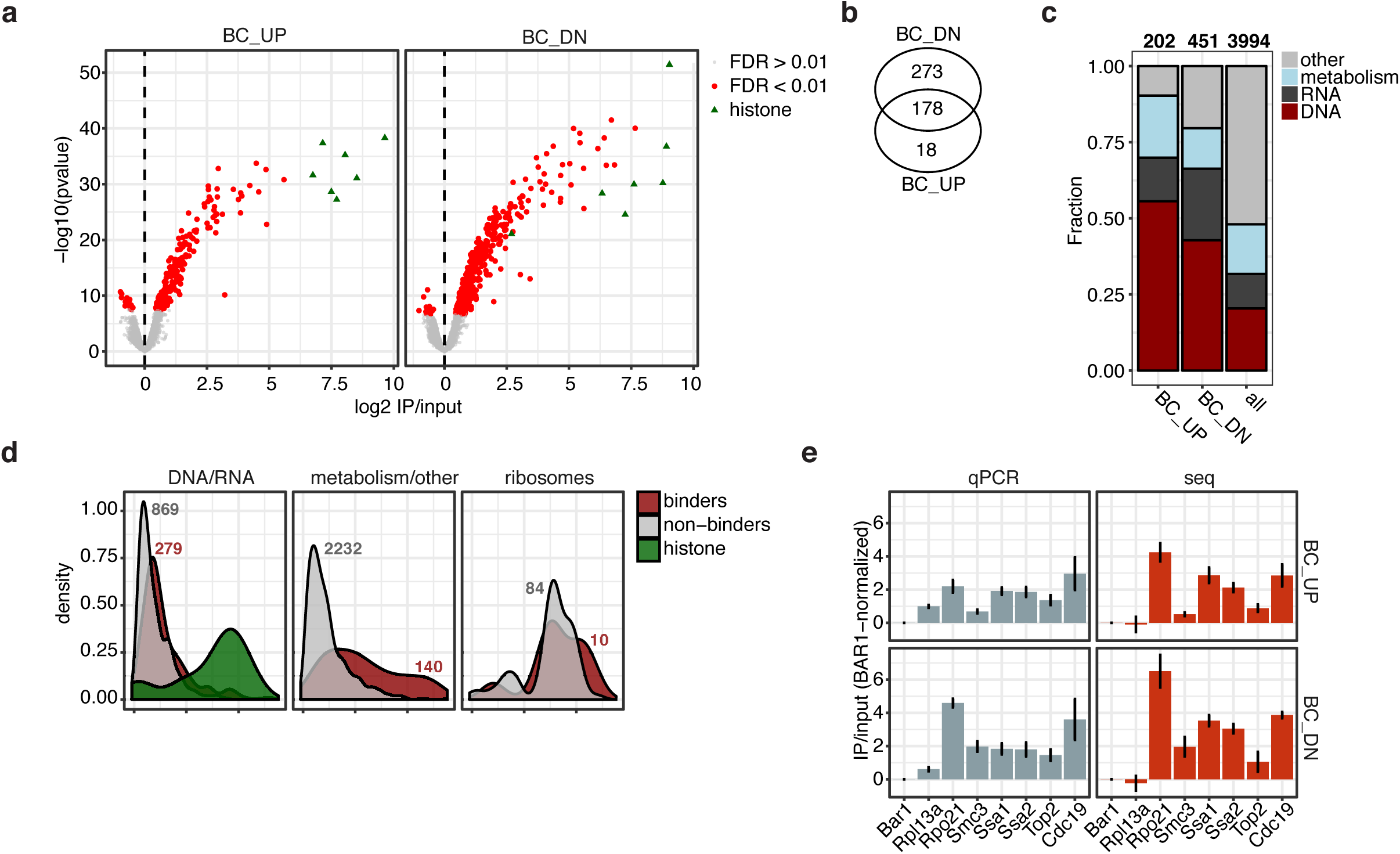
Identifying chromatin binders with DNA barcode counting. (A) Volcano plot showing the average log2 ChIP/input versus significance (p-value) calculated for 3,994 factors for BC_UP and 3,995 factors for BC_DN across 6 independent biological experiments. The red color indicates factors with a Benjamini-Hochberg adjusted p-value < 0.01. Green triangles represent the barcodes associated with the TAP-tagged histone proteins in the library. (B) Venn diagram indicating the number of factors that are significant (FC>0; FDR<0.01) at BC_UP, BC_DN, or both. (C) Bar plots showing the fraction of factors in four different categories that were determined by using GO annotations (see methods). The first two bars show the fractions in the group of binders (at either BC_UP or BC_DN). The right bar shows the fraction in the background set (the entire library). (D) Density plot of GFP abundance using data from the Cyclops database (http://cyclops.ccbr.utoronto.ca/). The binders were defined as before (FC>0;FDR<0.01), and the categories are similar as in C. The ribosome category was defined based on the CYC2008 database (http://wodaklab.org/cyc2008/). Histones are shown separately; they bind at the DNA and are highly abundant. The numbers indicate the size of each group. (E) ChIP-qPCR of selected TAP-tag strains from the Epi-Decoder library, with specific primers in proximity to BC_UP and BC_DN. Barl-TAP was used as a negative control because it is not expressed in these cells. To compare the barcode counts with ChIP-qPCR signal, the samples were normalized to the Bar1-TAP signal before calculating ChIP/input. The average of three biological replicates is shown; the error bars indicate the standard deviation. Rpl31a is a ribosomal subunit (frequently used in anchor away studies), Rpo21 is the largest subunit of RNA polymerase II, Smc3 is a cohesin subunit, Ssa1 and Ssa2 are HSP70 chaperones and 98% identical, Top2 is topoisomerase 2, and Cdc19/Pyk1 is pyruvate kinase.

### Different interactomes of the promoter and terminator regions

To determine whether Epi-Decoder can be used to identify locus-specific proteins, we next compared the significantly enriched factors for BC_UP to those of BC_DN. For this purpose, we plotted the binding scores for BC_UP versus BC_DN and color coded enriched factors according to the main chromatin functions or complexes expected at this locus (Fig. 3a). Strikingly, we observed that proteins within the same complex or process tend to have similar BC-specific binding scores and therefore tend to cluster together. This strongly suggests that Epi-Decoder does not just detect binding events but also provides quantitative information about protein occupancy.

**Figure 3.**
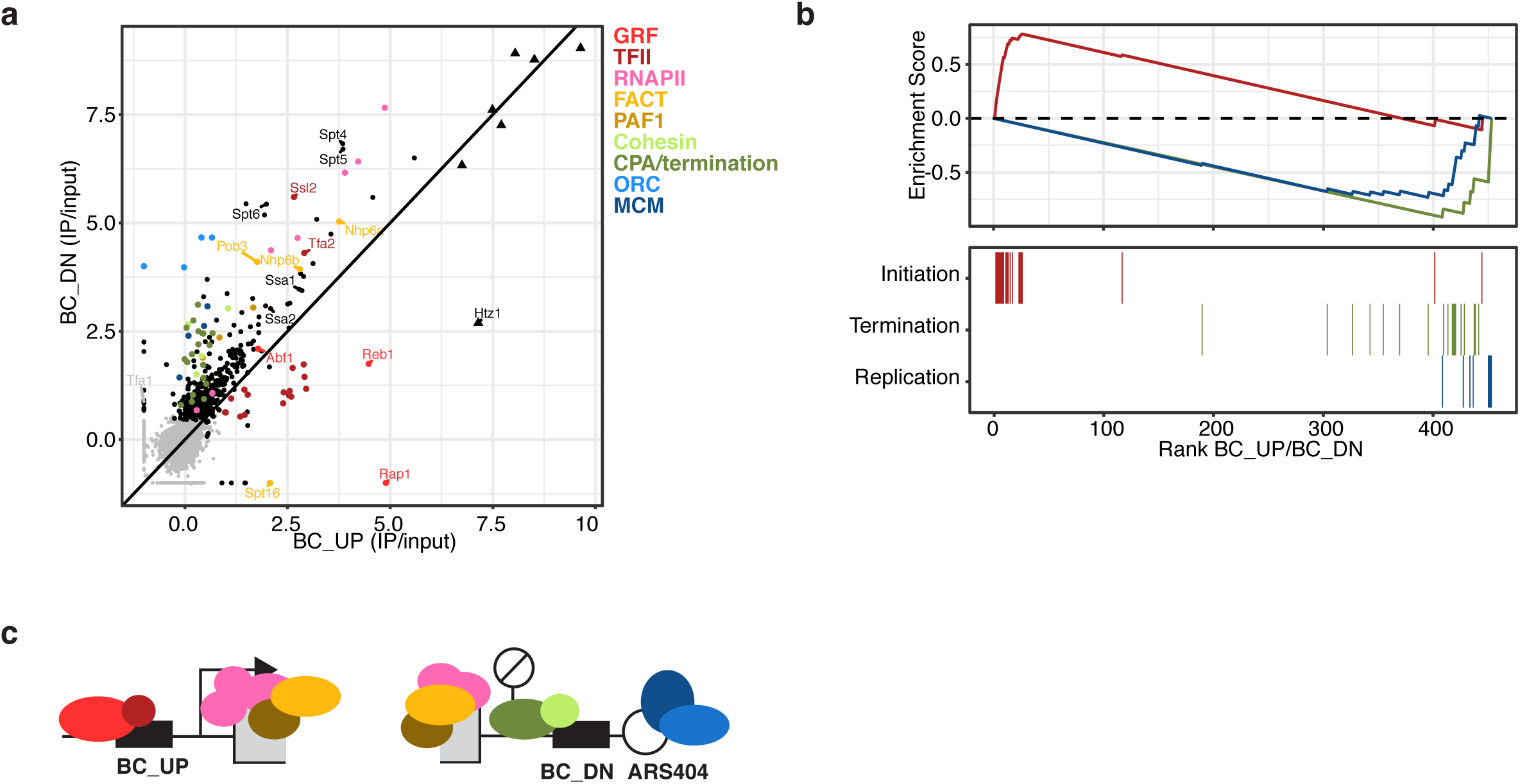
Capturing the chromatin interactome with Epi-Decoder. (A) Scatter plot of ChIP/input values, which can be interpreted as binding scores, of BC_UP versus BC_DN. For some factors, only the BC_UP or BC_DN value was available. In these cases, the missing value was set to −1 in order to still visualize the remaining value. Factors are colored based on known functions or complexes. Triangles represent histone proteins. The average values of six replicates are shown. (B) Enrichment plot for three categories: initiation (TFIID-E and general regulatory factors (GRFs)), termination (Cleavage and Polyadenylation and THO complex) and replication (ORC and MCM) factors. Binders were ranked based on the BC_UP/BC_DN ratio. The top part shows the running sum enrichment for each category. Initiation factors were significantly enriched at BC_UP (p=3.18^−3^), and termination and replication at BC_DN (p=2.03^−7^ and p=1.06^−5^). The bottom lines indicate the factors represented in the categories. (C) Illustration of the different protein complexes that Epi-Decoder identified at this barcoded HO locus.

Several factors and complexes were shared between the two locations. Besides histones, strongly-enriched factors at both barcodes include subunits of RNA polymerase II and several other factors and complexes with a well-known role in transcription elongation such as DSIF, Elf1, FACT, Spt6, and the PAF-C complex (Fig. 3a)^20-21^. However, BC_UP and BC_DN also showed substantial quantitative interactome differences reflecting their different functional states and suggesting that many proteins show locus-specific binding behavior (Fig. 3b). At BC_UP at the 5’ end of the gene, histone variant H2A.Z (*HTZ1*) was strongly enriched, which is in agreement with the enrichment of H2A.Z in promoter regions^22^. In addition, BC_UP showed enrichment for general regulatory factors Reb1 and Rap1. This is in agreement with the known Rap1 binding site in the *TEF1* promoter of the *KanMX* gene^23^ and the role of Reb1 (and Abf1) as a general chromatin organizer of regulatory regions^20^. BC_UP also showed enrichment for the basal transcription factors TFIID, -E, -F and –H, which form the pre-initiation complex together with RNA polymerase II^21,24^.

At the 3’ end, BC_DN showed enrichment for factors involved in transcription termination and mRNA cleavage and polyadenylation, an important step in mRNA 3’ end formation^25^. The Cohesin complex was also enriched at BC_DN. Finally, we observed strong binding of the evolutionary conserved ORC and MCM complexes at BC_DN, which is in agreement with the proximal origin of replication sequence at which ORC and then MCM are loaded to assemble the pre-replication complex^26-27^. Thus, in addition to common binders, Epi-Decoder revealed locus-specific enrichment of factors (Fig. 3c). This confirms that while the barcodes are in the same genomic region, they are in distinct functional contexts that can be separated by this approach even though they are only 1.5 kb apart. Interestingly, not all factors followed the expected distribution. Tfa2, the small subunit of the heterodimeric basal transcription factor TFIIE, and Ssl2, the dsDNA translocase subunit of the basal transcription factor TFIIH, were found at BC_UP but were also strongly enriched at BC_DN. 3’ binding of Tfa2 and Ssl2 has been observed at other genes as well^24,28^ but the significance remained uncertain. Since in Epi-Decoder all proteins have the same tag and are assessed simultaneously in a pool in a quantitative manner, the deviant binding pattern observed here cannot be explained by antibody issues or experimental and strain differences and therefore strongly suggests that Tfa2 and Ssl2 have special roles or positions within their complex or have non-canonical functions outside their complex.

### Conditional local proteomes: Chromatin rewiring in hydroxyurea

The chromatin interactome defined by Epi-Decoder confirmed that the barcodes flank an actively transcribed gene in close proximity to and transcribing towards an origin of replication that is licensed with the pre-replicative complex. This conformation is of special interest because upon firing of the origin, replication fork progression will require negotiation with the transcription machinery, potentially leading to collisions^29-33^. It has been proposed that in order to avoid collisions, RNA polymerase II is degraded at a subset of genes that are about to be replicated^34^. Since this recently described process of chromatin adaptation is still poorly understood, we used Epi-Decoder to determine in a comprehensive and unbiased manner how the barcoded HO locus interactome changes when cells are arrested in early S-phase in hydroxyurea (HU; Supplementary Fig. 3a), the condition in which RNA polymerase II degradation has been observed^34^. HU affects replication-fork progression by reducing the supply of dNTPs and by the generation of reactive oxygen species^35^.

We first confirmed previous observations^36^ that in HU early origins have fired but that the middle to late barcoded HO origin (ARS404) has not fired yet (Fig. 4a). Despite the absence of initiation of replication, Epi-Decoder revealed multiple changes in the chromatin proteome; 40 proteins showed a significantly different binding score in HU at BC_UP and 79 were significantly different at BC_DN and their altered abundance could not be explained by changes in protein levels (Supplementary Fig. 3b). Firstly, we observed lower mRNA levels of the barcoded *KanMX* gene compared to the *HphMX* gene, which expresses a different mRNA (Hph instead of Kan) from the same *AgTEF1* promoter but at a different genomic location (*HphMX* replaces the *CAN1* gene at chromosome V) and not in proximity to an origin of replication (Fig. 4b). This was accompanied by a reduction in occupancy of multiple RNA polymerase II subunits at the barcoded gene, extending the previous findings that RNA polymerase II is degraded at a subset of genes that have been or are about to be replicated (Fig. 4c). Secondly, the chromatin response was not restricted to RNA polymerase II because other general transcription proteins involved in capping, initiation, elongation and termination were also reduced, showing that the reduction of transcription involves a broad range of changes in the (co)transcriptional machinery (Fig. 4c). Thirdly, we observed additional changes that were not directly related to transcription but indicate topological alterations (Fig. 4c and Supplementary Fig. 3c). Topoisomerase II (Top2) binding was strongly increased at both ends of the *KanMX* gene in HU. Top2 is the main enzyme releasing topological stress during S phase^37^ and targeted to nucleosome free DNA during replication stress^38^. Pds5, the cohesin maintenance factor was also increased at BC_DN, and a closer inspection of the cohesin complex indicates that three of the four cohesin subunits in the library showed moderately increased occupancy as well, suggesting stabilization of cohesin binding in this region (Supplementary Table 2). Finally, all the subunits of the ORC complex in the library showed decreased binding at BC_DN, while occupancy of the MCM complex was unaltered. ORC proteins have been suggested to remain bound to origins throughout the yeast cell cycle^39-40^, possibly being negatively influenced by MCM proteins in G1^41^. The direct and quantitative comparison of all origin-proximal factors by Epi-Decoder suggest that the interaction of ORC proteins with the origin is compromised in S-phase prior to firing but that this cannot be explained by increased MCM protein occupancy. Therefore, our results show that ORC subunits interact more dynamically with chromatin throughout the yeast cell-cycle than expected but are in agreement with the behavior of ORC in other organisms^39,42^. How lower ORC binding influences origin firing and subsequent fork progression of this locus remains unknown. Interestingly, recent *in vitro* reconstitution experiments demonstrated that the MCM complex is stably bound to DNA once assembled and also competent for replication, even after removal of ORC proteins, suggesting that origin firing perse may not be affected^43^.

**Figure 4.**
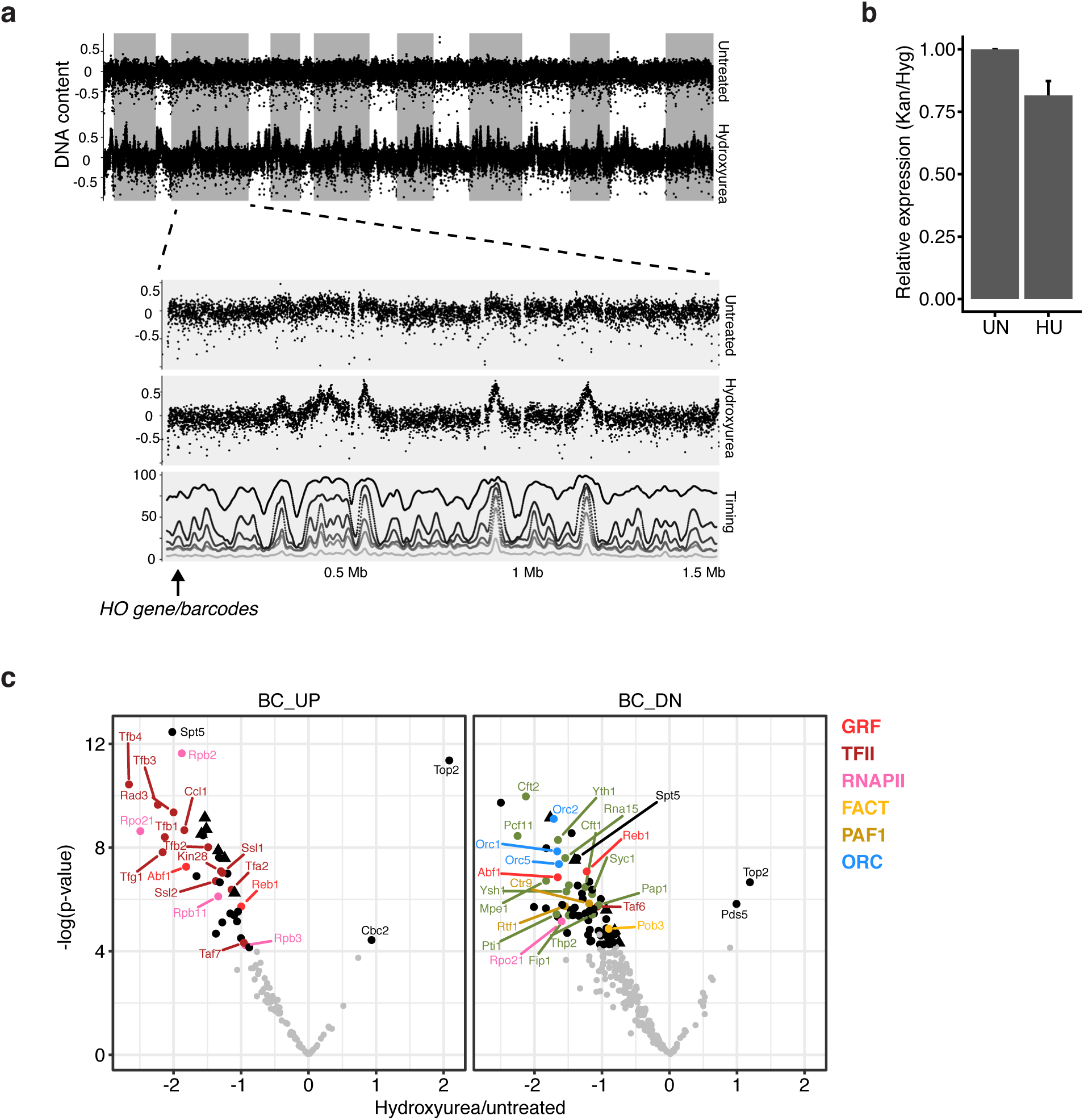
Chromatin rewiring upon hydroxyurea treatment. (A) DNA regions replicated in the HU arrest were determined by DNA copy number assessment. The DNA coverage in bins of 200 bp is plotted across the genome for untreated and hydroxyurea treated samples. The lower panel shows a zoom-in of chromosome IV, which contains the barcoded locus on the left arm (indicated by the arrow). Replication timing in the absence of HU was obtained from Alvino *et al*^36^. The percentage of replicated DNA is plotted for each bin at 10, 12.5, 15, 17.5, 25 and 40 minutes after release from a G1 arrest, as indicated by the shades of grey. (B) RT-qPCR shows relative mRNA expression of KanMX/HphMX in untreated and hydroxyurea-treated samples. The untreated samples were set to 1. The average of three biological replicates is shown; error bars indicate standard deviation. (C) Volcano plot showing the ratio of binding in hydroxyurea/untreated for factors that were significantly enriched in either one or both of the conditions. The colored dots are factors with significantly different binding scores (FDR < 0.05)

In summary, Epi-Decoder uncovered condition-dependent composition of a local chromatin-proteome. At a transcribed gene next to a licensed origin, arrest in HU led to tuning down of transcription by a general reduction of transcription protein binding, which is accompanied by increased occupancy of topology factors, and altered stoichiometry of replication proteins.

## Discussion

Strategies aimed at identifying proteins bound at specific genomic loci frequently involve affinity-capture combined with mass spectrometry^5-6^. Affinity handles generally involve engineering the locus by introducing binding sites for a tagged protein, capturing a native locus by using oligo-capture, or targeting inactive tagged Cas9 proteins. Important progress has been made for multi-copy DNA loci such as repetitive DNA elements^7-8^. Recently, oligo-capture^44^ and CRISPR-Cas9 technology^45^ have been applied to identify proteins at single copy genomic loci. However, the problem of assessing a unique locus in a comprehensive and quantitative manner by mass spectrometry has not been solved and many challenges remain, a major one being the high degree of purification of the locus of interest while obtaining sufficient material for good coverage in MS analysis^6^. Epi-Decoder provides a powerful and orthogonal strategy in which the very challenging proteomics problem of decoding the chromatin proteome of a single genomic locus is addressed by DNA sequencing. We demonstrate that Epi-Decoder enables an unbiased, comprehensive and quantitative analysis of protein occupancy of a single locus in budding yeast.

In addition to known and expected core chromatin proteins, we observed significant binding scores for many proteins that do not have canonical DNA-related functions. Among the unexpected factors are several metabolic enzymes, oxidative-stress response factors, the yeast ubiquitin activating enzyme, chaperones, proteasome subunits, and RNA-processing factors (Supplementary Table 1). Some of these unexpected factors are highly expressed; their binding score may reflect non-specific binding. On the other hand, not all unexpected interactors are highly expressed or show equal binding scores. In addition, for a substantial number of the identified metabolic enzymes (e.g. glyceraldehyde 3-phosphate dehydrogenase/Tdh3, pyruvate kinase/Cdc19, homocitrate synthase/Lys20,Lys21) and protein chaperones (e.g. the HSP70 chaperones Ssa1 and Ssa2), there is evidence that they have active roles in transcription and DNA repair^46-50^. Furthermore, the binding scores of some metabolic enzymes changed upon treatment with HU (Supplementary Table 2), indicating that their interaction with chromatin is not constitutive. Indeed, there is a growing body of evidence that enzymes in the cytoplasm might have moonlighting functions in the nucleus, for example by local supply of co-factors of histone modifying enzymes^19,46,51-54^. Alternatively, the interaction with chromatin could somehow influence the activity of metabolic enzymes. The barcoded *HO* locus data we describe here provides a rich resource for further unraveling the biological meaning of non-canonical chromatin interactions.

Epi-Decoder is more than a discovery tool. It offers the possibility to compare two loci or to analyze one locus under different physiological or genetic conditions in a systematic and quantitative manner. Since all proteins are examined in a pooled fashion and using the same affinity handle, differences in purification and cell backgrounds can be excluded and binding scores can be directly compared. This is exemplified by the basal transcription factors Tfa2 and Ssl2, which are quantitative outliers compared to their complex members (TFIIE and TFIIH, respectively), suggesting non-canonical functions. Another example is the chromatin rewiring we observed upon treatment with HU, which uncovers large scale quantitative changes in interactions of transcription and replication proteins, as well as other factors. We note that Epi-Decoder generates independent binding scores for proteins with (nearly) identical protein sequences but encoded by different genes, such as the histone proteins or other protein paralogs such as Ssa1 and Ssa2, which are 98% identical. This can provide extra information about the differential use of related proteins that cannot easily or not be distinguished by mass spectrometry methods. Protein isoforms that arise through post-translational modification cannot be distinguished by Epi-Decoder.

The Epi-Decoder strategy will be applicable to a broad range of fundamental epigenetic questions. A few requirements need to be met for successful application. First, Epi-Decoder requires a library of tagged proteins. For budding yeast, several tagged protein libraries are already available in addition to the carboxy-terminal TAP-tag library used here^55-57^. This will allow for extending the current Epi-Decoder analyses to a nearly complete coverage of the proteome as well as to alternative (amino-terminal) tags, together enabling an unprecedented deep analysis of chromatin proteomes. Importantly, tagged protein libraries are also becoming available for many other organisms^58-62^. A second requirement for Epi-Decoder applications is the integration of DNA barcodes proximal to the locus of interest. Given their small size, DNA barcodes generally only minimally disrupt a locus but this needs to be verified for the locus of interest, as is the case for strategies involving targeting affinity handles by Cas9 or other approaches. Fortunately, genomic barcoding has come within reach for many research questions due to the availability emerging genome engineering strategies, in yeast and other organisms^63-66^. For example, in budding yeast, CRIPSR-Cas9 tools provide the means to integrate barcodes from simple oligonucleotide-derived repair templates with very high efficiency and precision without the need for a selectable marker. This greatly facilitates creating a library of barcoded strains with minimal disruption of the locus of interest.

In summary, Epi-Decoder is powerful and versatile strategy for decoding the protein interactome of a single genomic locus. We expect that the Epi-Decoder strategy and derivatives thereof will enable the decoding of dynamic proteomes of different loci in different organisms and will be of high value for addressing a broad range of important chromatin-biology questions in future applications.

## Methods

### Yeast strains and library construction

Yeast strains used in this study are listed in Supplementary Table 3. Library manipulations on solid media were performed using synthetic genetic array (SGA) technology^12^ combined with robotics using a RoToR machine (Singer Instruments, Watchet, UK). A collection of barcoded Tandem Affinity Purification (TAP)-tagged strains (library NKI4217) was generated as follows. In order to cross the two libraries, the TAP-tag collection^15^ was made compatible for SGA and converted to mating type α. This was done by crossing the TAP library with NKI4212, which was derived from strain Y8205 by replacing the STE2pr-Sp_his5 cassette at the *CAN1* locus by *HphMX* to enable selection during SGA for the TAP-His3MX6 alleles by using histidine prototrophy. This resulted in the *MATα* TAP collection (NKI4214) divided over 4 different 1536-plates that were each mated with the set of 1140 barcoder *MATa* strains^14^. Diploids were obtained by G418 and Hygromycin double selection on rich media and transferred to sporulation media. After sporulation, the combination library was obtained by selecting twice for *MATα* haploids and then twice for *MATα* haploids containing a TAP-tag, the barcoded KanMX cassette and the HphMX marker. The barcodes and TAP tags were verified for a few selected strains. Five extra barcoded control strains (TAP-tagged versions of Hht2, Htb2, Rpl13a, Ste2 and Bar1) were generated by using the respective clones from the NKI4214 library and introducing unused barcodes from *MATa* haploid gene knockout library (Open Biosystems, Huntsville, AL) at the *HO* locus of these strains.

### Media and growth conditions

Yeast media was prepared as described previously^12,67^. For screening in untreated conditions, yeast strains were grown in YEPD (1% yeast extract, 2% bacto peptone, and 2% glucose) in log phase. S phase arrest was achieved by adding to log phase cells (OD660 of 0.4) in YEPD one volume of YEPD + 360 mM HU to achieve 180 mM HU final concentration and cells were harvested after 2 hours. The arrest was verified by flow cytometry.

### Epi-Decoder (TAG-ChIP-Barcode-Seq)

The NKI4217 libraries were grown on YEPD plates overnight and the colonies of each plate were pooled together in liquid culture. The cultures were grown until log phase (OD660 of ~0.4) and cross-linked for 20 minutes with 1/10th of the volume of freshly prepared Fix Solution (1% formaldehyde, 50 mM Hepes-KOH, pH 7.5, 100 mM NaCl, 1 mM EDTA) and subsequent quenched for 5 minutes with Glycine (125 mM final concentration). Cells were washed once in cold TBS with 0.2 mM PMSF and the pellet was frozen at −80°C. Cells from frozen pellets of ~1.5x10^9^ cells were lysed by bead beating in 200 μL breaking buffer (100 mM Tris pH 7.9, 20% glycerol, protease inhibitor cocktail EDTA-free) with Zirconia/silica beads. The lysate was washed twice in 1 ml FA buffer (50 mM HEPES-KOH pH 7.5, 140 mM NaCl, 1 mM EDTA, 1% Triton X-100, 0.1% Na-deoxycholate, protease inhibitor cocktail EDTA-free) and sonicated using the Bioruptor PICO (Diagenode) for 10 minutes at 30 s intervals. Chromatin was cleared by centrifugation for 5 minutes at 4 °C at 4000 rpm. One hundred microliter chromatin was used as input material. For ChIP, IgG Sepharose 6 Fast Flow beads (GE healthcare) were washed 3 times with PBSB (PBS containing 5 mg/mL BSA), and incubated with 1 ml chromatin for 6 hours on a turning wheel at 4 °C. Samples were washed twice in FA buffer, twice in high salt FA buffer (500 mM NaCl), twice in RIPA buffer (10 mM Tris pH 8, 250 mM LiCl, 0.5% NP-40, 0.5% Na-deoxycholate, 1 mM EDTA) and once with TE buffer (10 mM Tris pH8, 1 mM EDTA). With each wash step, the Sepharose beads were spun for 2 minutes at 3000 rpm at 4 °C. IP samples were eluted for 10 minutes at 65°C in 100 μL elution buffer (50 mM Tris pH 8, 10 mM EDTA, 1% SDS). IP and input samples were digested with 0.5 μL RNase A (10 mg/mL) and 10 μL ProtK (10 mg/mL) in 70 μL TE for 1 hour at 50 °C and subsequently kept overnight at 65°C to reverse crosslinks. DNA was purified using the QIAquick PCR purification kit (Qiagen). BC_UP and BC_DN were amplified separately with specific primers (Supplementary Table 4), mixed in an equimolar fashion and purified from an agarose gel with a size selection of 100-150 bp. The purified DNA was sequenced (single read, >50bp) on a HiSeq2500/MiSeq platform (Illumina, San Diego, CA), using a mix of custom sequencing primers (Supplementary Table 4).

### Barcode counting

Barcodes were extracted from the sequencing reads by using the Perl script extracting Counting and Linking to Barcode References (XCALIBR). The code and detailed descriptions of the functions are available at ttps://github.com/NKI-GCF/xcalibr. Briefly, XCALIBR locates the constant regions (U2 and D2) in each amplicon and reports the 6 bp upstream for the index and 20 bp downstream for the barcode. The resulting table contains counts for each barcode-index combination. Counts below 10 were removed from this table before further pre-processing and filtering steps.

### Barcode data pre-processing and filtering

Input (IN) and immunoprecipitated DNA (IP) of each plate was amplified separately with a unique index. Even though we aimed to mix each sample equimolarly, plate-specific differences in counts could still occur. This was corrected by normalizing each plate by its median. The counts table was log2 transformed and barcode-index combinations were matched with the ORF names. Factors with low input counts were removed since these were likely to be missing from the library or the barcode failed to amplify due to technical reasons. We manually validated several factors for the presence of a TAP tag by PCR with primers in the specific ORF and the TAP tag. This revealed that ORC4, MCM3, MCM7 and RAD6 were not properly tagged; we therefore removed these strains for further analysis. The final set contained information on 3,994 BC_UP and 3,955 BC_DN barcode clones. We noticed that the dynamic range was slightly different between biological replicates. To overcome this problem, we performed quantile-normalization for the replicates of BC_UP and BC_DN separately. This method is used to generate similar distributions by first ranking each replicate based on the counts and then replacing the counts of each rank by the mean count of that rank.

### Barcode statistical analysis

Factors with IP counts that were significantly enriched over input were identified by using the Limma R/Bioconductor software package^68^. P-values were adjusted for multiple testing by converting them to false discovery rates using the Benjamini-Hochberg procedure. Factors with a positive fold-change and an FDR<0.01 were selected as significantly enriched. For untreated conditions, six independent biological samples were used. The running sum scores for the enrichment plot (Fig. 3b) were calculated with the gseaScores function from the HTSanalyzeR package. For identifying differential binders upon HU treatment, three biological samples were used for treatment and no treatment. Here, we only considered factors that were significant binders in at least one of the conditions (FC>0 and FDR<0.05). Limma was used to select factors with significantly different IP counts (FDR<0.05). Barcode counts were compared with protein abundance data measured by GFP intensity. This data was obtained from the CYCLoPs database^69^.

### GO slim process enrichment

GO slim process terms are condensed versions of the full GO ontology^70-71^. GO slim process terms were downloaded from the Saccharomyces Genome Database^72^ and enrichment analysis was performed by using the fisher.test function in R with option alternative=“greater”. We manually assigned the following categories based on GO slim terms listed here. DNA binding: DNA-templated transcription, initiation, DNA-templated transcription, elongation, DNA-templated transcription, termination, transcription from RNA polymerase I promoter, transcription from RNA, polymerase II promoter, chromatin organization, histone modification, DNA replication, DNA recombination, DNA repair, cellular response to DNA damage stimulus, regulation of DNA metabolic process, nuclear transport and chromosome segregation. RNA binding or processing: mRNA processing, rRNA processing, tRNA processing, RNA modification, RNA splicing and RNA catabolic process. Metabolism: carbohydrate metabolic process, cofactor metabolic process, nucleobase-containing small molecule metabolic process, monocarboxylic acid metabolic process, cellular amino acid metabolic process, generation of precursor metabolites and energy, oligosaccharide metabolic process and lipid metabolic process.

### ChIP-qPCR

ChIP experiments were performed similar to Tag-ChIP-Barcode-Seq, but with 20 μl bed volume of IgG Sepharose 6 Fast Flow beads and 200 μl chromatin. Factors were selected such that they reflect different classes of binders, in addition to negative controls Bar1 (not expressed in these cells) and Rpl13A (a ribosomal subunit). Quantitative PCR was performed on the purified DNA with SYBR green master mix (Applied Biosystems or Roche) or SensiFAST SYBR master mix (Bioline) according to the manufacturer's protocol and analyzed on LightCycler 480 II (Roche). The binding was analyzed with specific primers in close proximity to the BC_UP and BC_DN (Supplementary Table 4). Each sample was measured in two technical duplicates in the qPCR and the average value of these two was taken as one value when combining biological replicates. In order to compare the ChIP-qPCR and BC-seq results in a quantitative manner, the negative control (Bar1) was used to normalize the raw values.

### Flow Cytometry

Flow cytometry samples were prepared to monitor cell cycle progression and verify S phase arrest by HU. 1×10^7^Cells were collected and fixed with 70% ethanol and stored at −20 °C. Flow cytometry was performed as previously described^73^, after staining DNA with Sytox green (Molecular Probes). Flow cytometry measurements were taken on a FACSCalibur with CellQuest software (Becton Dickinson) and further analyzed with FlowJo software (Treestar).

### Genomic DNA isolation and sequencing

Two independent strains were selected for genomic DNA isolation, the strains with tagged versions of Bar1 and Rpo21. Genomic DNA was isolated as described previously^74^. Briefly, 2x10^7^ cells were spun down and resuspended in 200 μl of 2% Triton X-100, 1% SDS, 100 mM NaCl, 10 mM Tris pH 8.0, 1 mM EDTA. Then, 200 μl phenol-chloroform-isoamyl alcohol (25:24:1) was added and cells were lysed by bead beating with 300 μl Zirconia/silica beads. 200 μl TE was added, the cells were spun for 5 minutes at max speed and the aqueous layer was transferred to a new tube. The pellet was washed in 1 ml 100% ethanol and re-suspended in 400 μl TE. To precipitate the DNA 10 ~14 M ammonium acetate and 1 ml 100% ethanol was added and the tube was inverted to mix contents and spun 2 minutes at 13000 rpm. The pellet was air dried and suspend in 50 μl TE. The gDNA was then purified by sending through an Isolate II genomic DNA kit (Bioline). 20 μl of ethanol precipitated gDNA was resuspended in 180 μl lysis buffer GL and 1 μl RNaseA (10 mg/ml) and kept at room temperature for 20 minutes before proceeding with step 3 of the manufacturers protocol. DNA was eluted in 50 μl elution buffer G. The gDNA was sheared with a Covaris E220 ultrasonicator (Covaris) to obtain fragments of 150-200bp and sequenced on the HiSeq2000 platform (Illumina, San Diego, CA).

### Genomic DNA isolation and sequencing data analysis

Single-end reads of 65 bp were aligned to the *S. cerevisiae* reference genome version R64 (UCSC SacCer3) by using BWA^75^. The BAM files with aligned reads were filtered for mapq 37 and converted to bedgraph with bins of 250 bp by using deepTools^76^. By using custom R scripts, the region of unstable transcript XUT_12F-188 was removed because it contains repetitive sequences. The average coverage in each 250 bp bin was plotted across the entire genome by using custom R scripts. The replication timing in the absence of HU was obtained from a density transfer experiment^36^, and downloaded from Table S1 in that study. This table contains the percentage of the genome that had become hydrid in density (%HL DNA) in cells collected at different time points^36^. The plots were generated for both samples separately and showed similar patterns.

### Reverse Transcription

RNA was isolated using the RNeasy Mini Kit (QIAGEN) using the protocol for Yeast cells, with a few modifications. Briefly, 2x10^7^ cells were spun down (5 minutes at 3000 rpm) and pellets were dissolved in 600 μL cold RLT buffer. Cells were broken by bead beating with 400 μL Zirconia/silica beads and debris was separated by centrifuging 2 minutes at 13000 rpm. The supernatant (~350 μL) was collected and mixed with 1V (~350 μL) 70% EtOH and transported to RNeasy columns. Following the buffer RW1 and buffer RPE wash steps, RNA was eluted in 50uL elution buffer. Eluted RNA was treated with DNase I (QIAGEN) to remove genomic DNA. Next, cDNA was prepared using superscript II reverse transcriptase (Invitrogen). RT-PCR was performed with the primers in Supplementary Table 4. Each sample was measured in two technical duplicates and the average value of these two was taken as one value when combining biological replicates.

## Acknowledgements

The authors thank the RHPC facility of the Netherlands Cancer Institute for providing computational resources, and Hanneke Vlaming for advice on barcode sequencing. This work was supported by the Netherlands Organisation for Scientific Research (NWO-VICI-016.130.627). The funders had no role in study design, data collection and interpretation, or the decision to submit the work for publication. We thank Fabio Brocco, Dominique Morais, Ila van Kruijsbergen, Thom Molenaar, Eliza Mari Kwesi-Maliepaard, Oscar Aparicio, Maarten Altelaar, Bas van Steensel, Tineke Lenstra, and Haico van Attikum for valuable suggestions.

## Author contributions

FvL and KV conceived the project. DPL, TK and KV designed experiments and performed the screens. TK performed bioinformatics analyses. DPL performed validation experiments and analyzed the results. TvW assisted with validation experiments. FvL supervised the project. FvL, TK, and DPL wrote the paper.

## Abbreviations

BC: Barcode
ChIP: Chromatin Immunoprecipitation
SGA: Synthetic Genetic Array technology
SGD: Saccharomyces Genome Database
TAP: Tandem Affinity Purification
HU: Hydroxyurea

---

**Supplementary Figure 1.**
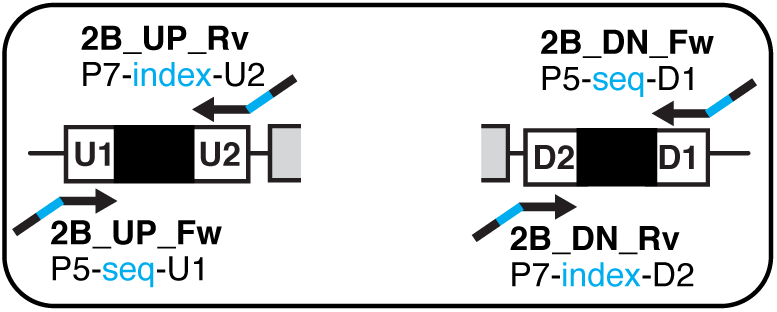
Schematic overview of the primers used to amplify the barcodes, which are flanked by constant regions U1, U2, D1 and D2. The forward primers introduce the Illumina P5 sequence and extra nucleotides for annealing of the 5' end of the custom sequencing primers. The reverse primers introduce the Illumina P7 sequence as well as a 6 bp index

**Supplementary Figure 2.**
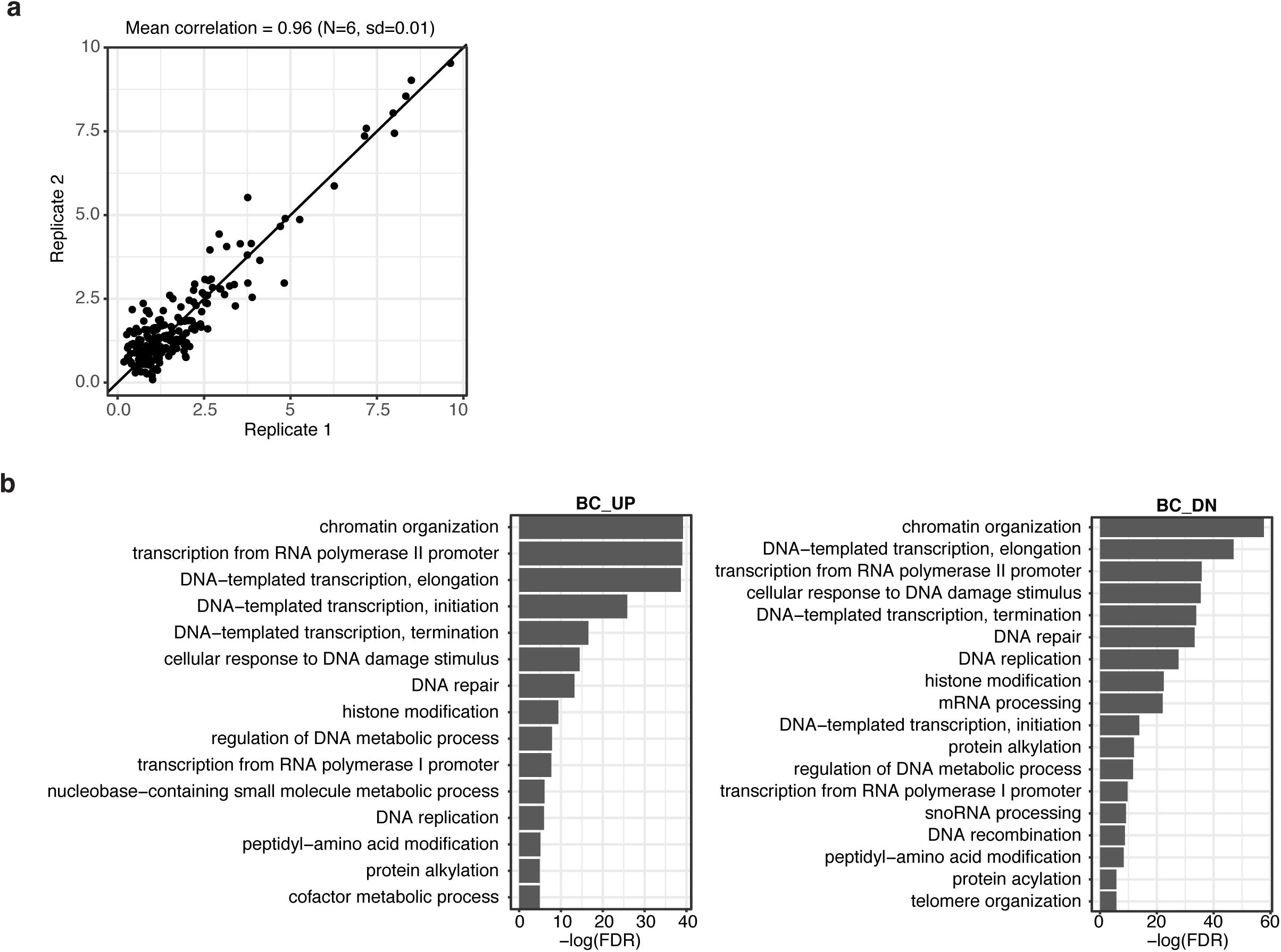
(A) Scatter plot showing the normalized barcode counts for the binders of biological replicates. Two samples were randomly chosen for a representative figure. The mean correlation and standard deviation was calculated for all six replicates. (B) Bar plot showing the GO slim terms (process) enriched at BC_UP and BC_DN. Terms with a Fisher Exact p-value < 0.01 were considered to be enriched. The x-axis shows the number of factors associated with each term, and the non-transparent color indicates the proportion of binders.

**Supplementary Figure 3.**
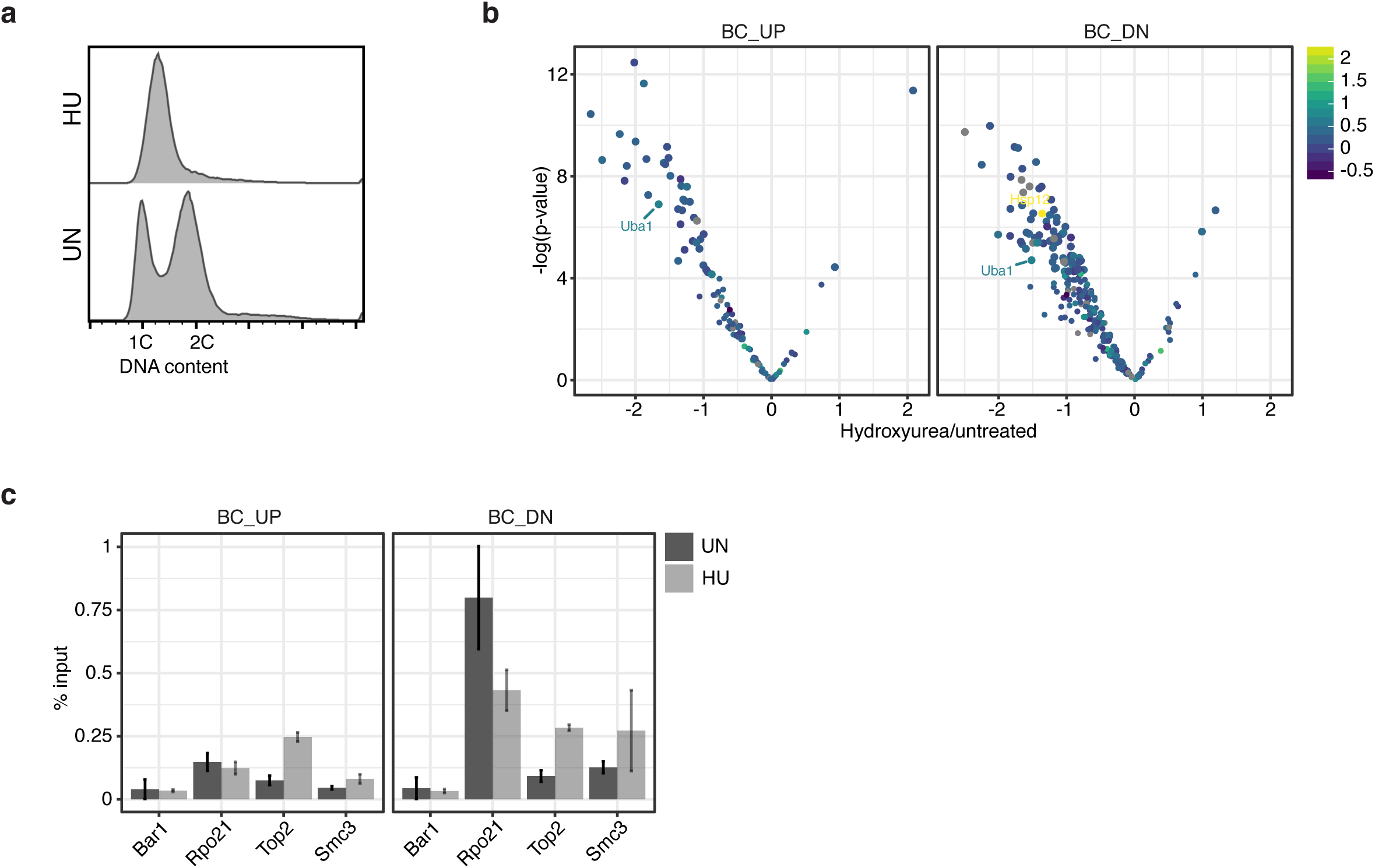
(A) Flow cytometry analysis showing the DNA content of cells with (HU) and without hydroxyurea (UN) treatment (180 mM for 2 hours). (B) Volcano plot similar to Figure 4b, but here the dots are colored based on their protein abundance changes in HU based on data from^77^. (C) ChIP-qPCR for selected strains, with specific primers in close proximity to BC_UP and BC_DN. Bar1-TAP was used as a negative control because it is not expressed in these cells. To compare the barcode counts with ChIP-qPCR signal, the samples were normalized by the Bar1-TAP signal before calculating ChIP/input. The average of three biological replicates is shown; the error bars indicate the standard deviation.

**Supplementary Table 1**

List of strains in the Epi-decoder library and their associated binding scores. For each TAP-tagged strain, the corresponding barcode enrichment (average log2 fold-change (logFC) of IP/In) and Benjamini-Hochberg corrected p-value (FDR) are provided for both BC_UP and BC_DN. The GO_cat column contains the general categories based on GO slim process terms. The complex column contains several well-known protein complexes that bind DNA.

**Supplementary Table 2**

List of binders in either untreated or hydroxyurea treated cells and their associated binding difference. This table contains the significant binders in either untreated or hydroxyurea treatment. For each strain, the average log2 fold-change (logFC) of hydroxyurea/untreated and their associated Benjamini-Hochberg corrected p-value (FDR) are provided for both BC_UP and BC_DN. The complex column contains several well-known protein complexes that bind DNA.

**Supplementary Table 3.**
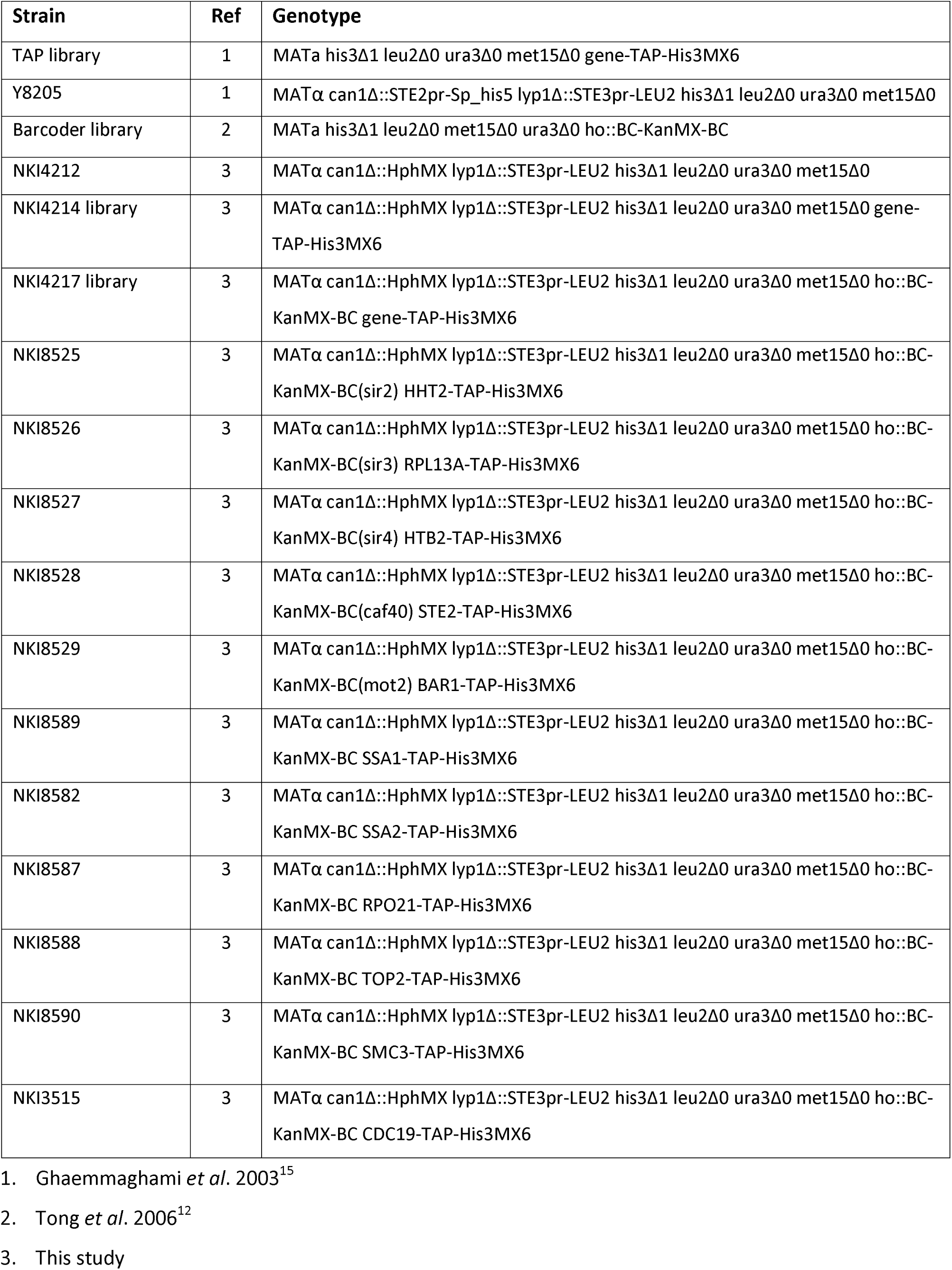
List of yeast strains used in this study.

| Strain | Ref | Genotype |
| --- | --- | --- |
| TAP library | 1 | MATa his3Δ1 leu2Δ0 ura3Δ0 met15Δ0 gene-TAP-His3MX6 |
| Y8205 | 1 | MATα can1Δ::STE2pr-Sp_his5 lyp1Δ::STE3pr-LEU2 his3Δ1 leu2Δ0 ura3Δ0 met15Δ0 |
| Barcoder library | 2 | MATa his3Δ1 leu2Δ0 met15Δ0 ura3Δ0 ho::BC-KanMX-BC |
| NKI4212 | 3 | MATα can1Δ::HphMX lyp1Δ::STE3pr-LEU2 his3Δ1 leu2Δ0 ura3Δ0 met15Δ0 |
| NKI4214 library | 3 | MATα can1Δ::HphMX lyp1Δ::STE3pr-LEU2 his3Δ1 leu2Δ0 ura3Δ0 met15Δ0 gene-TAP-His3MX6 |
| NKI4217 library | 3 | MATα can1Δ::HphMX lyp1Δ::STE3pr-LEU2 his3Δ1 leu2Δ0 ura3Δ0 met15Δ0 ho::BC-KanMX-BC gene-TAP-His3MX6 |
| NKI8525 | 3 | MATα can1Δ::HphMX lyp1Δ::STE3pr-LEU2 his3Δ1 leu2Δ0 ura3Δ0 met15Δ0 ho::BC-KanMX-BC(sir2) HHT2-TAP-His3MX6 |
| NKI8526 | 3 | MATα can1Δ::HphMX lyp1Δ::STE3pr-LEU2 his3Δ1 leu2Δ0 ura3Δ0 met15Δ0 ho::BC-KanMX-BC(sir3) RPL13A-TAP-His3MX6 |
| NKI8527 | 3 | MATα can1Δ::HphMX lyp1Δ::STE3pr-LEU2 his3Δ1 leu2Δ0 ura3Δ0 met15Δ0 ho::BC-KanMX-BC(sir4) HTB2-TAP-His3MX6 |
| NKI8528 | 3 | MATα can1Δ::HphMX lyp1Δ::STE3pr-LEU2 his3Δ1 leu2Δ0 ura3Δ0 met15Δ0 ho::BC-KanMX-BC(caf40) STE2-TAP-His3MX6 |
| NKI8529 | 3 | MATα can1Δ::HphMX lyp1Δ::STE3pr-LEU2 his3Δ1 leu2Δ0 ura3Δ0 met15Δ0 ho::BC-KanMX-BC(mot2) BAR1-TAP-His3MX6 |
| NKI8589 | 3 | MATα can1Δ::HphMX lyp1Δ::STE3pr-LEU2 his3Δ1 leu2Δ0 ura3Δ0 met15Δ0 ho::BC-KanMX-BC SSA1-TAP-His3MX6 |
| NKI8582 | 3 | MATα can1Δ::HphMX lyp1Δ::STE3pr-LEU2 his3Δ1 leu2Δ0 ura3Δ0 met15Δ0 ho::BC-KanMX-BC SSA2-TAP-His3MX6 |
| NKI8587 | 3 | MATα can1Δ::HphMX lyp1Δ::STE3pr-LEU2 his3Δ1 leu2Δ0 ura3Δ0 met15Δ0 ho::BC-KanMX-BC RPO21-TAP-His3MX6 |
| NKI8588 | 3 | MATα can1Δ::HphMX lyp1Δ::STE3pr-LEU2 his3Δ1 leu2Δ0 ura3Δ0 met15Δ0 ho::BC-KanMX-BC TOP2-TAP-His3MX6 |
| NKI8590 | 3 | MATα can1Δ::HphMX lyp1Δ::STE3pr-LEU2 his3Δ1 leu2Δ0 ura3Δ0 met15Δ0 ho::BC-KanMX-BC SMC3-TAP-His3MX6 |
| NKI3515 | 3 | MATα can1Δ::HphMX lyp1Δ::STE3pr-LEU2 his3Δ1 leu2Δ0 ura3Δ0 met15Δ0 ho::BC-KanMX-BC CDC19-TAP-His3MX6 |

1. Ghaemmaghami *et al.* 2003<sup>15</sup> 2. Tong *et al.* 2006<sup>12</sup> 3. This study

**Supplementary Table 4.**
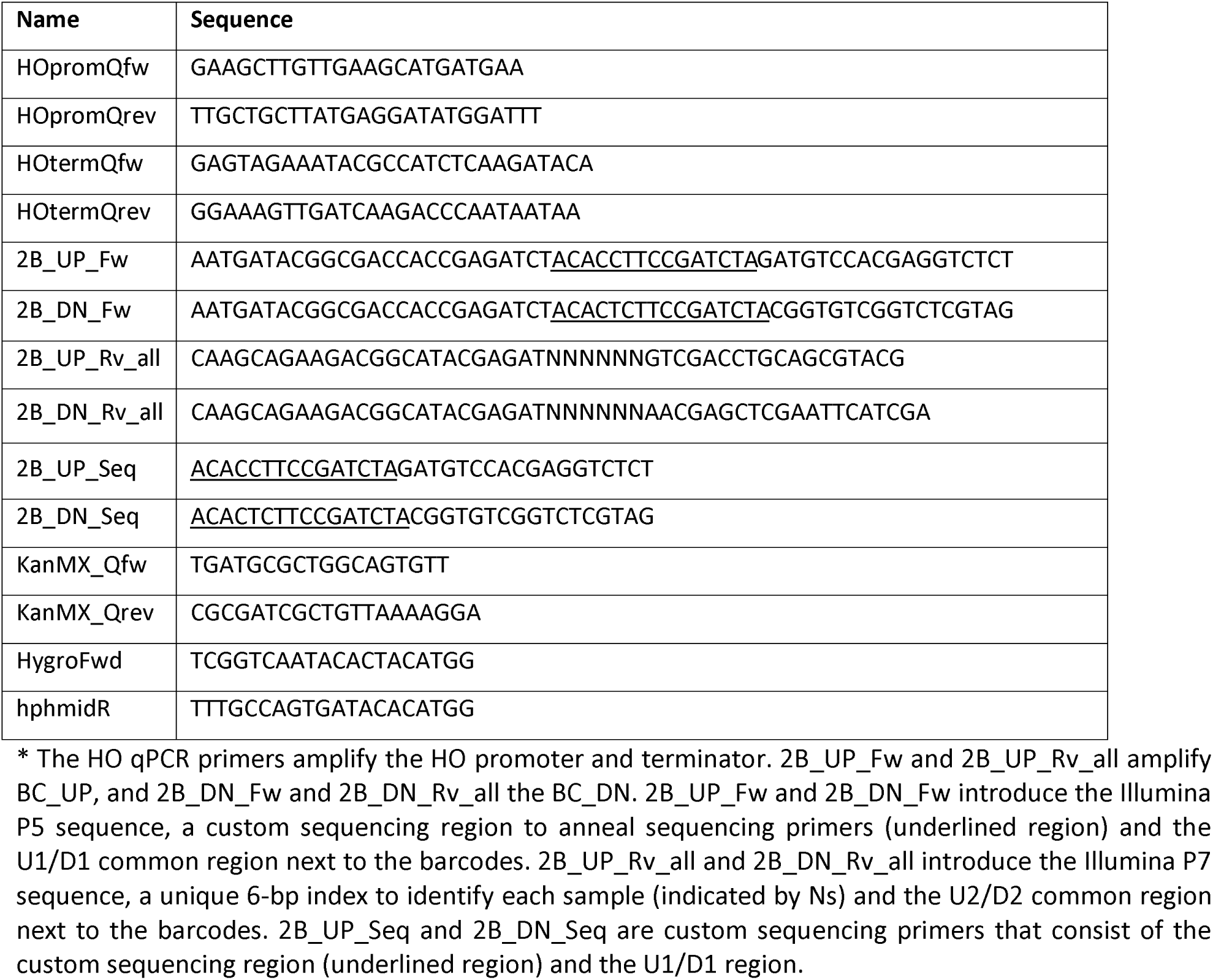
List of the primers used in this study*.

